# Human demographic history impacts genetic risk prediction across diverse populations

**DOI:** 10.1101/070797

**Authors:** Alicia R. Martin, Christopher R. Gignoux, Raymond K. Walters, Genevieve L. Wojcik, Benjamin M. Neale, Simon Gravel, Mark J. Daly, Carlos D. Bustamante, Eimear E. Kenny

## Abstract

The vast majority of genome-wide association studies are performed in Europeans, and their transferability to other populations is dependent on many factors (e.g. linkage disequilibrium, allele frequencies, genetic architecture). As medical genomics studies become increasingly large and diverse, gaining insights into population history and consequently the transferability of disease risk measurement is critical. Here, we disentangle recent population history in the widely-used 1000 Genomes Project reference panel, with an emphasis on populations underrepresented in medical studies. To examine the transferability of single-ancestry GWAS, we used published summary statistics to calculate polygenic risk scores for six well-studied traits and diseases. We identified directional inconsistencies in all scores; for example, height is predicted to decrease with genetic distance from Europeans, despite robust anthropological evidence that West Africans are as tall as Europeans on average. To gain deeper quantitative insights into GWAS transferability, we developed a complex trait coalescent-based simulation framework considering effects of polygenicity, causal allele frequency divergence, and heritability. As expected, correlations between true and inferred risk were typically highest in the population from which summary statistics were derived. We demonstrated that scores inferred from European GWAS were biased by genetic drift in other populations even when choosing the same causal variants, and that biases in any direction were possible and unpredictable. This work cautions that summarizing findings from large-scale GWAS may have limited portability to other populations using standard approaches, and highlights the need for generalized risk prediction methods and the inclusion of more diverse individuals in medical genomics.

## Introduction

The majority of genome-wide association studies (GWAS) have been performed in populations of European descent^1–4^. An open question in medical genomics is the degree to which these results transfer to new populations. GWAS have yielded tens of thousands of common genetic variants significantly associated with human medical and evolutionary phenotypes, most of which have replicated in other ethnic groups^5–8^. However, GWAS are optimally powered to discover common variant associations, and the European bias in GWAS results in associated SNPs with higher minor allele frequencies on average compared to other populations. The predictive power of GWAS findings in non-Europeans are therefore limited by population differences in allele frequencies and linkage disequilibrium structure.

As GWAS sample sizes grow to hundreds of thousands of samples, they also become better powered to detect rare variant associations^9–11^. Large-scale sequencing studies have demonstrated that rare variants show stronger geographic clustering than common variants^12–14^. Rare, disease-associated variants are therefore expected to track with recent population demography and/or be population restricted^13,15–17^. As the next era of GWAS expands to evaluate the disease-associated role of rare variants, it is not only scientifically imperative to include multi-ethnic populations, it is also likely that such studies will encounter increasing genetic heterogeneity in very large study populations. A comprehensive understanding of the genetic diversity and demographic history of multi-ethnic populations is critical for appropriate applications of GWAS, and ultimately for ensuring that genetics does not contribute to or enhance health disparities^4^.

The most recent release of the 1000 Genomes Project (phase 3) provides one of the largest global reference panels of whole genome sequencing data, enabling a broad survey of human genetic variation^18^. The depth and breadth of diversity queried facilitates a deep understanding of the evolutionary forces (e.g. selection and drift) shaping existing genetic variation in present-day populations that contribute to adaptation and disease^19–25^. Studies of admixed populations have been particularly fruitful in identifying genetic adaptations and risk for diseases that are stratified across diverged ancestral origins^26–34^. Admixture patterns became especially complex during the peopling of the Americas, with extensive recent admixture spanning multiple continents. Processes shaping structure in these admixed populations include sex-biased migration and admixture, isolation-by-distance, differential drift in mainland versus island populations, and variable admixture timing^13,35,36^.

Standard GWAS strategies approach population structure as a nuisance factor. A typical step-wise procedure first detects dimensions of global population structure in each individual, using principal component analysis (PCA) or other methods^37–40^, and often excludes “outlier” individuals from the analysis and/or corrects for inflation arising from population structure in the statistical model for association. Such strategies reduce false positives in test statistics, but can also reduce power for association in heterogeneous populations, and are less likely to work for rare variant association^41–44^. Recent methodological advances have leveraged patterns of global and local ancestry for improved association power^30,45,46^, fine-mapping^47^ and genome assembly^48^. At the same time, population genetic studies have demonstrated the presence of fine-scale sub-continental structure in the African, Native American, and European components of populations from the Americas^49–52^. If trait-associated variants follow the same patterns of demography, then we expect that modeling sub-continental ancestry may enable their improved detection in admixed populations.

In this study, we explore the impact of population diversity on the landscape of variation underlying human traits. We infer demographic history for the global populations in the 1000 Genomes Project, focusing particularly on admixed populations from the Americas, which are under represented in medical genetic studies^4^. We disentangle local ancestry to infer the ancestral origins of these populations. We link this work to ongoing efforts to improve study design and disease variant discovery by quantifying biases in clinical databases and GWAS in diverse and admixed populations. These biases have a striking impact on genetic risk prediction; for example, a previous study calculated polygenic risk scores for schizophrenia in East Asians and Africans based on GWAS summary statistics derived from a European cohort, and found that prediction accuracy was reduced by more than 50% in non-European populations^53^. To disentangle the role of demography on polygenic risk prediction derived from single-ancestry GWAS, we designed a novel coalescent-based simulation framework reflecting modern human population history and show that polygenic risk scores derived from European GWAS are biased when applied to diverged populations. Specifically, we identify reduced variance in risk prediction with increasing divergence from Europe reflecting decreased overall variance explained, and demonstrate that an enrichment of low frequency risk and high frequency protective alleles contribute to an overall protective shift in European inferred risk on average across traits. Our results highlight the need for the inclusion of more diverse populations in GWAS as well as genetic risk prediction methods improving transferability across populations.

## Material and Methods

### Ancestry deconvolution

We used the phased haplotypes from the 1000 Genomes consortium. We phased reference haplotypes from 43 Native American samples from^54^ inferred to have > 0.99 Native ancestry in ADMIXTURE using SHAPEIT2 (v2.r778)^55^, then merged the haplotypes using scripts made publicly available. These combined phased haplotypes were used as input to the PopPhased version of RFMix v1.5.4^56^ with the following flags: -w 0.2, -e 1, -n 5, –use-reference-panels-in-EM, –forward-backward. The node size of 5 was selected to reduce bias in random forests resulting from unbalanced reference panel sizes (AFR panel N=504, EUR panel N=503, and NAT panel N=43). We used the default minimum window size of 0.2 cM to enable model comparisons with previously inferred models using *Tracts*^*57*^. We used 1 EM iteration to improve the local ancestry calls without substantially increasing computational complexity. We used the reference panel in the EM to take better advantage of the Native American ancestry tracts from the Hispanic/Latinos in the EM given the small NAT reference panel. We set the LWK, MSL, GWD, YRI, and ESN as reference African populations, the CEU, GBR, FIN, IBS, and TSI as reference European populations, and the samples from Mao et al^54^ with inferred > 0.99 Native ancestry as reference Native American populations, as previously^58^.

### Ancestry-specific PCA

We performed ancestry-specific PCA, as described in^35^. The resulting matrix is not necessarily orthogonalized, so we subsequently performed singular value decomposition in python 2.7 using numpy. There were a small number of major outliers, as seen previously^35^. There was one outlier (ASW individual NA20314) when analyzing the African tracts, which was expected as this individual has no African ancestry. There were 8 outliers (PUR HG00731, PUR HG00732, ACB HG01880, ACB HG01882, PEL HG01944, ACB HG02497, ASW NA20320, ASW NA20321) when analyzing the European tracts. Some of these individuals had minimal European ancestry, had South or East Asian ancestry misclassified as European ancestry resulting from a limited 3-way ancestry reference panel, or were unexpected outliers. As described in the PCAmask manual, a handful of major outliers sometimes occur. As AS-PCA is an iterative procedure, we therefore removed the major outliers for each sub-continental analysis and orthogonalized the matrix on this subset.

### Tracts

The RFMix output was collapsed into haploid bed files, and “UNK” or unknown ancestry was assigned where the posterior probability of a given ancestry was < 0.90. These collapsed haploid tracts were used to infer admixture timings, quantities, and proportions for the ACB and PEL (new to phase 3) using *Tracts*^*57*^. Because the ACB have a very small proportion of Native American ancestry, we fit three 2-way models of admixture, including one model of single- and two models of double-pulse admixture events, using *Tracts*. In both of the double-pulse admixture models, the model includes an early mixture of African and European ancestry followed by another later pulse of either European or African ancestry. We randomized starting parameters and fit each model 100 times and compared the log-likelihoods of the model fits. The single-pulse and double-pulse model with a second wave of African admixture provided the best fits and reached similar log-likelihoods, with the latter showing a slight improvement in fit.

We next assessed the fit of 9 different models in *Tracts* for the PEL^57^, including several two-pulse and three-pulse models. Ordering the populations as NAT, EUR, and AFR, we tested the following models: ppp_ppp, ppp_pxp, ppp_xxp, ppx_xxp, ppx_xxp_ppx, ppx_xxp_pxx, ppx_xxp_pxp, ppx_xxp_xpx, and ppx_xxp_xxp, where the order of each letter corresponds with the order of populations given above, an underscore indicates a distinct migration event with the first event corresponding with the most generations before present, p corresponding with a pulse of the ordered ancestries, and x corresponding with no input from the ordered ancestries. We tested all 9 models preliminarily 3 times, and for all models that converged and were within the top 3 models, we subsequently fit each model with 100 starting parameters randomizations.

### Imputation accuracy

Imputation accuracy was calculated using a leave-one-out internal validation approach. Two array designs were compared for this analysis: Illumina OmniExpress and Affymetrix Axiom World Array LAT. Sites from these array designs were subset from chromosome 9 of the 1000 Genomes Project Phase 3 release for admixed populations. After fixing these sites, each individual was imputed using the rest of the dataset as a reference panel.

Overall imputation accuracy was binned by minor allele frequency (0.5-1%, 1-2%, 2-3%, 3-4%, 4-5%, 5-10%, 10-20%, 20-30%, 30-40%, 40-50%) comparing the genotyped true alleles to the imputed dosages. A second round of analyses stratified the imputation by local ancestry diplotype, which was estimated as described earlier. Within each ancestral diplotype (AFR_AFR, AFR_NAT, AFR_EUR, EUR_EUR, EUR_NAT, NAT_NAT), imputation accuracy was again estimated within MAF bins.

### Empirical polygenic risk score inferences

To compute polygenic risk scores in the 1000 Genomes samples using summary statistics from previous GWAS, we first filtered to biallelic SNPs and removed ambiguous AT/GC SNPs from the integrated 1000 Genome call set. To get relatively independent associations taking LD into account when multiple significant p-value associations are in the same region in a GWAS, we performed LD clumping in plink (-clump) for all variants with MAF ≥ 0.01^59^, which uses a greedy algorithm ordering SNPs by p-value, then selectively removes SNPs within close proximity and LD in ascending p-value order (i.e. starting with the most significant SNP). As a population cohort with similar LD patterns to the study sets, we used European 1000 Genomes samples (CEU, GBR, FIN, IBS, and TSI). To compute the polygenic risk scores, we considered all SNPs with p-values ≤ 1e-2 in the GWAS study, a window size of 250 kb, and an R^2^ threshold of 0.5 in Europeans to group SNPs. After obtaining the top clumped signals, we computed scores using the –score flag in plink.

### Polygenic risk score simulations

We simulated genotypes in a coalescent framework with msprime v0.4.0^60^ for chromosome 20 incorporating a recombination map of GRCh37 and an assumed mutation rate of 2e-8 mutations / (base pair * generation). We used a demographic model previously inferred using 1000 Genomes sequencing data^13^ to simulate individuals that reflect European, East Asian, and African population histories. We focus on these populations as the demography has previously been modeled and this avoids the challenges of simulating the geographically heterogeneous^52^ and sex-biased process of admixture in the Americas^61^. To imitate a GWAS with European sample bias and evaluate polygenic risk scores in other populations, we simulated 200,000 European individuals, 200,000 East Asian, and 200,000 African individuals. Next, we assigned “true” causal effect sizes to *m* evenly spaced alleles. Specifically, we randomly assigned effect sizes as:

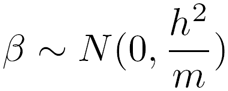

where the normal distribution is specified by the mean and standard deviation (as in python’s numpy package). For all other non-causal sites, the effect size is zero. We then define *X* as:

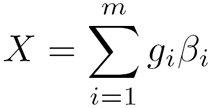

where the *g*_*i*_ are the genotype states (i.e. 0, 1, or 2). To handle varying allele frequencies, potential weak LD between causal sites, ensure a neutral model with random true polygenic risks with respect to allele frequencies, and to obtain the total desired variance, we normalize *X* as:

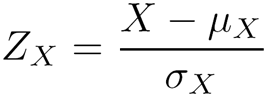

We then compute the true polygenic risk score, as:

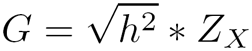

such that the total variance of the scores is *h*^2^. We also simulated environmental noise and standardize to ensure equal variance between normalized genetic and environmental effects before, defining the environmental effect *E* as:

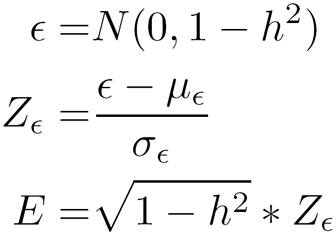

such that the total variance of the environmental effect is *1 – h*^*2*^. We then define the total liability as:

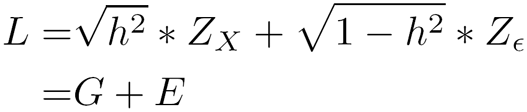

We assigned 10,000 European individuals at the most extreme end of the liability threshold “case” status assuming a prevalence of 5%. We randomly assigned 10,000 different European individuals “control” status. We ran a GWAS with these 10,000 European cases and 10,000 European controls, computing Fisher’s exact test for all sites with MAF > 0.01. As before for empirical polygenic risk score calculations from real GWAS summary statistics, we clumped these SNPs into LD blocks for all sites with p ≤ 1e-2, R^2^ ≥ 0.5 in Europeans, and within a window size of 250 kb. We used these SNPs to compute inferred polygenic risk scores as before, summing the product of the log odds ratio and genotype for the true polygenic risk in a cohort of 10,000 simulated European, African, and East Asian individuals (all not included in the simulated GWAS). We compared the true versus inferred polygenic risk scores for these individuals across varying complexities (*m* = 200, 500, 1000) and heritabilites (*h*^2^ = 0.33, 0.50, 0.67).

### Results

#### Genetic diversity within and between populations in the Americas

We first assessed the overall diversity at the global and sub-continental level of the 1000 Genomes Project (phase 3) populations^18^ using a likelihood model via ADMIXTURE^62^ and PCA^63^ (**Figure S1** and **Figure S2**). The six populations from the Americas demonstrate considerable continental admixture, with genetic ancestry primarily from Europe, Africa, and the Americas, recapitulating previously observed population structure^18^. To quantify continental genetic diversity in these populations, we repeated the analysis using YRI, CEU, and NAT samples^54^ as reference panels (population labels and abbreviations in **Table S1**). We observed widely varying continental admixture contributions in the six populations from the Americas at K=3 (Figure 1A and **Table S2**). For example, when compared to the ASW, the ACB have a higher proportion of African ancestry (μ = 0.88, 95% CI = [0.87-0.89] versus μ = 0.76, 95% CI = [0.73-0.78]; two-sided t-test p=3.0e-13) and a smaller proportion of EUR and NAT ancestry. The PEL have more NAT ancestry than all of the other AMR populations (μ = 0.77, 95% CI = [0.75-0.80] versus CLM: μ = 0.26, 95% CI = [0.24, 0.27], p=2.9e-95; PUR: μ = 0.13, 95% CI = [0.12, 0.13], p=4.8e-93; and MXL: μ = 0.47, 95% CI = [0.43, 0.50], p=1.7e-28) ascertained in 1000 Genomes.

**Figure 1.**
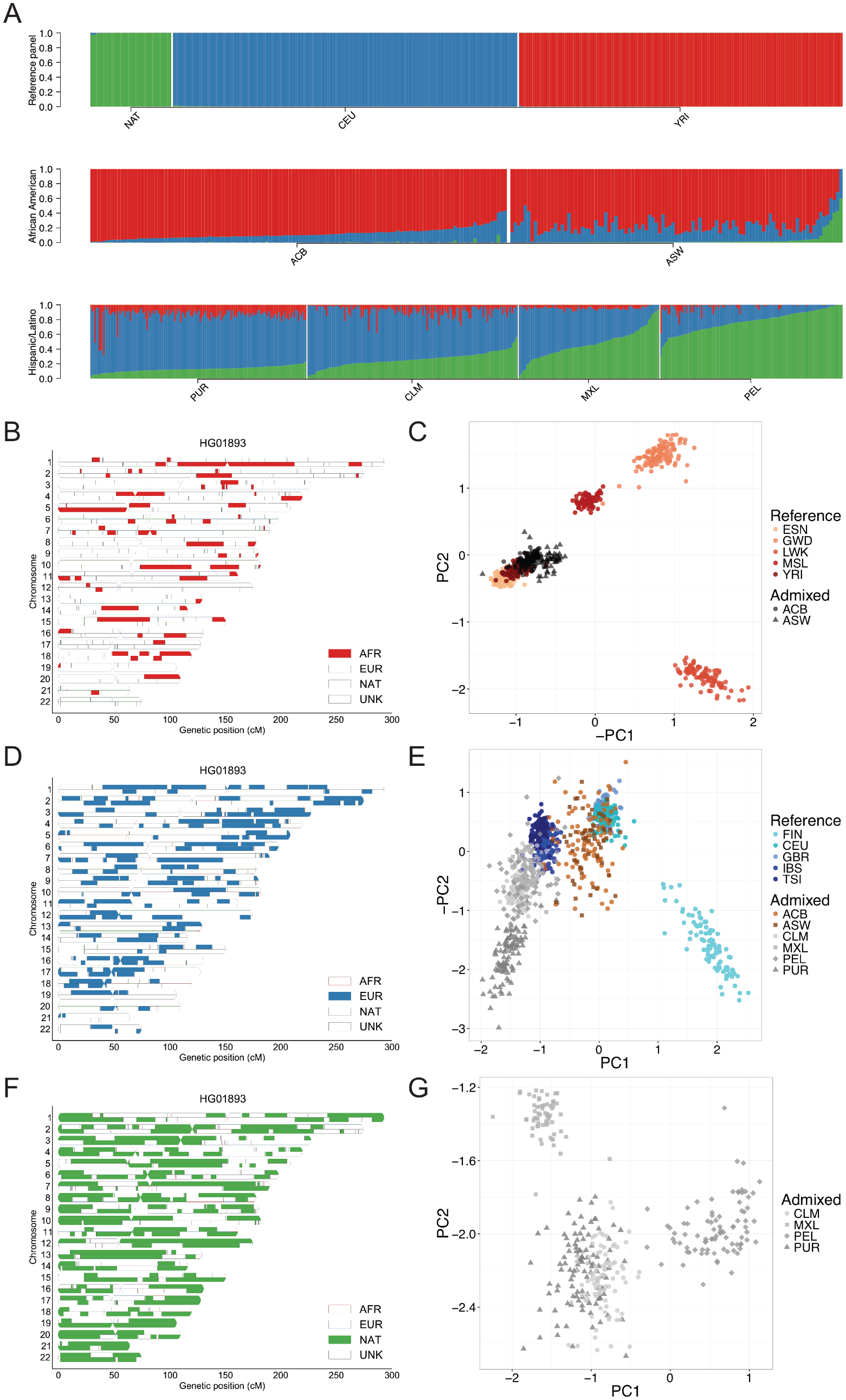
Sub-continental diversity and origins of African, European, and Native American components of recently admixed Americas populations. A) ADMIXTURE analysis at K=3 focusing on admixed Americas samples, with the NAT^54^, CEU, and YRI as reference populations. B,D,F) Local ancestry karyograms for representative PEL individual HG01893 with B) African, D) European, and F) Native American components shown. C,E,G) Ancestry-specific PCA applied to admixed haploid genomes as well as ancestrally homogeneous continental reference populations from 1000 Genomes (where possible) for C) African tracts, E) European tracts, and G) Native American tracts. A small number of admixed samples that constituted major outliers from the ancestry-specific PCA analysis were removed, including C) 1 ASW sample (NA20314) and E) 8 samples, including 3 ACB, 2 ASW, 1 PEL, and 2 PUR samples.

We explored the origin of the subcontinental-level ancestry from recently admixed individuals by identifying local ancestry tracts^29,35,56,64^ (Methods, **Figure S3**). As proxy source populations for the recent admixture, we used EUR and AFR continental samples from the 1000 Genomes Project as well as NAT samples genotyped previously^54^. Concordance between global ancestry estimates inferred using ADMIXTURE at K=5 and RFMix was typically high (Pearson’s correlation ≥ 98%, see **Figure S4**). Using *Tracts*^*57*^, we modeled the length distribution of the AFR, EUR, and NAT tracts to infer that admixing began ~12 and ~8 generations ago in the PEL and ACB populations, respectively (**Figure S5**), consistent with previous estimates from other populations from the Americas^49,57,65^.

We further investigated the subcontinental ancestry of admixed populations from the Americas one ancestry at a time using a version of PCA modified to handle highly masked data (ancestry-specific or AS-PCA) as implemented in PCAmask^66^. Example ancestry tracts in a PEL individual subset to AFR, EUR, and NAT components are shown in Figure 1B, D, and F, respectively. Consistent with previous observations, the inferred European tracts in Hispanic/Latino populations most closely resemble southern European IBS and TSI populations with some additional drift^35^ (Figure 1E). The

European tracts of the PUR are more differentiated compared to the CLM, MXL, and PEL populations, consistent with sex bias (**Figure S6 and Table S3**) and excess drift from founder effects in this island population^35^. In contrast to the southern European tracts from the Hispanic/Latino populations, the African descent populations in the Americas have European admixture that more closely resembles the northwestern CEU and GBR European populations. The clusters are less distinct, owing to lower overall fractions of European ancestry, however the European components of the Hispanic/Latino and African American populations are significantly different (Wilcoxan rank sum test p=2.4e-60).

The ability to localize aggregated ancestral genomic tracts enables insights into the evolutionary origins of admixed populations. To disentangle whether the considerable Native American ancestry in the ASW individuals arose from recent admixture with Hispanic/Latino individuals or recent admixture with indigenous Native American populations, we queried the European tracts. We find that the European tracts of all ASW individuals with considerable Native American ancestry are well within the ASW cluster and project closer in Euclidean distance with AS-PC1 and AS-PC2 to northwestern Europe than the European tracts from Hispanic/Latino samples (p=1.15e-3), providing support for the latter hypothesis and providing regional nuance to previous findings^49^.

We also investigated the African origin of the admixed AFR/AMR populations (ACB and ASW), as well as the Native American origin of the Hispanic/Latino populations (CLM, MXL, PEL, and PUR). The African tracts of ancestry from the AFR/AMR populations project closer to the YRI and ESN of Nigeria than the GWD, MSL, and LWK populations (Figure 1C). This is consistent with slave records and previous genome-wide analyses of African Americans indicating that most sharing occurred in West and West-Central Africa^67–69^. There are subtle differences between the African origins of the ACB and ASW populations (e.g. difference in distance from YRI on AS-PC1 and AS-PC2 p=6.4e-6), likely due either to mild island founder effects in the ACB samples or differences in African source populations for enslaved Africans who remained in Barbados versus those who were brought to the US. The Native tracts of ancestry from the AMR populations first separate the southernmost PEL populations from the CLM, MXL, and PUR on AS-PC1, then separate the northernmost MXL from the CLM and PUR on AS-PC2, consistent with a north-south cline of divergence among indigenous Native American ancestry (Figure 1G). ^35,70^

#### Impact of continental and sub-continental diversity on disease variant mapping

To investigate the role of ancestry in phenotype interpretation from genetic data, we assessed diversity across populations and local ancestries for recently admixed populations across the whole genome and sites from two reference databases: the GWAS catalog and ClinVar pathogenic and likely pathogenic sites. We recapitulate results showing that there is less variation across the genome (both genome-wide and on the Affymetrix 6.0 GWAS array sites used in local ancestry calling) in out-of-Africa versus African populations, but that GWAS variants are more polymorphic in European and Hispanic/Latino populations (**Figure S7A-B, Figure S8**A-B). We use a normalized measure of the minor allele frequency, an indicator of the amount of diversity captured in a population, to obtain a background coverage of each population, as done previously (e.g. Figure S4 from phase 3 of the 1000 Genomes Project^18^). We show that the Affymetrix 6.0 array has a slight European bias (**Figure S5**A and **Figure S6**A). We compared the site frequency spectrum of variants across the genome versus at GWAS catalog sites, and identify elevated allele frequencies at GWAS catalog loci, particularly in populations with more European ancestry (e.g. the EUR, AMR, and SAS super populations, **Figure S5**C-D). We further compared heterozygosity (estimated here as 2pg) and the site frequency spectrum in recently admixed populations across diploid and haploid local ancestry tracts, respectively. Sites in the GWAS catalog and ClinVar are more and less common than genome-wide variants, respectively (Figure 2). Whereas heterozygosity across the whole genome is highest in African ancestry tracts, it is consistently the greatest in European ancestry tracts across these databases (Figure 2 and **Figure S8**C-D), reflecting a strong bias towards European study participants^1–4,18,71^. These results highlight imbalances in genome interpretability across local ancestry tracts in recently admixed populations and the utility of analyzing these variants jointly with these ancestry tracts over genome-wide ancestry estimates alone.

**Figure 2.**
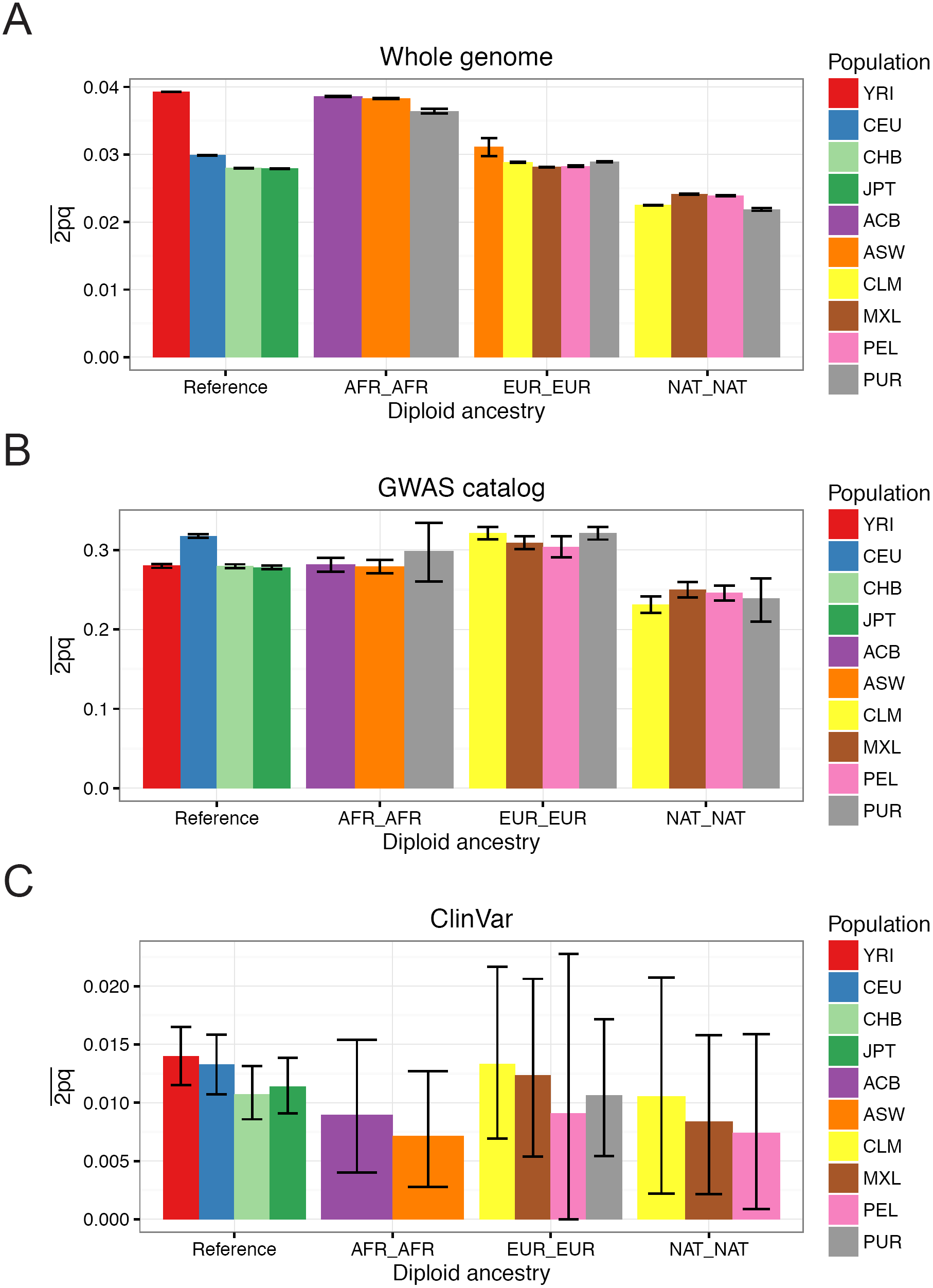
Heterozygosity (estimated here as 2pq) in admixed populations stratified by diploid local ancestry in A) the whole genome, B) sites from the GWAS catalog, and C) sites from ClinVar classified as “pathogenic” or “likely pathogenic.” The mean and 95% confidence intervals were calculatated by bootstrapping 1000 times. Populations not shown in a given panel have too few diploid ancestry tracts overlapping sites to calculate heterozygosity.

We also assessed imputation accuracy across the 3-way admixed populations from the Americas (CLM, MXL, PEL, PUR) for two arrays: the Illumina OmniExpress and the Affymetrix Axiom World Array LAT. Imputation accuracy was estimated as the correlation (r^2^) between the original genotypes and the imputed dosages. For both array designs, imputation accuracy across all minor allele frequency (MAF) bins was highest for populations with the largest proportion of European ancestry (PUR) and lowest for populations with the largest proportion of Native American ancestry (PEL, **Figure S9**A-B). We also stratified imputation accuracy by local ancestry tract diplotype within the Americas. Consistently, tracts with at least one Native American ancestry tract had lower imputation accuracy when compared to tracts with only European and/or African ancestry (Figure 3 and **Figure S10**).

**Figure 3.**
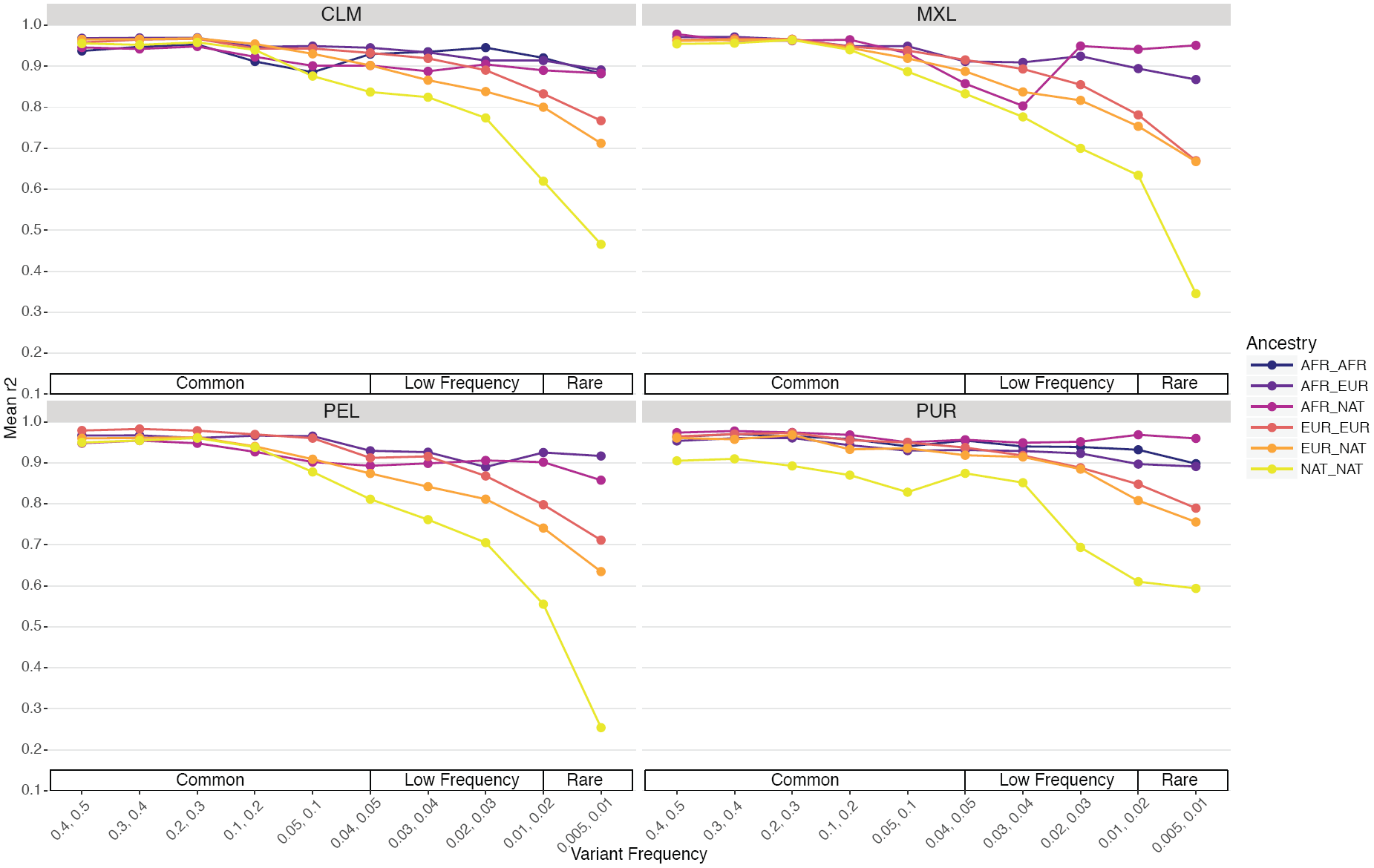
Imputation accuracy by population assessed using a leave-on-out strategy, stratified by diploid local ancestry on chromosome 9 for the Illumina OmniExpress genotyping array.

#### Transferability of GWAS findings across populations

To quantify the transferability of European-biased genetic studies to other populations, we next used published GWAS summary statistics to infer polygenic risk scores^72^ across populations for well-studied traits, including height^9^, waist-hip ratio^73^, schizophrenia^10^, type II diabetes^74,75^, and asthma^76^ (Figure 4A-D, **Figure S11,** Methods). Most of these summary statistics are derived from studies with primarily European cohorts, although GWAS of type II diabetes have been performed in both European-specific cohorts as well as across multi-ethnic cohorts. We identify clear directional inconsistencies in these score inferences. For example, although the height summary statistics show the expected cline of southern/northern cline of increasing European height (FIN, CEU, and GBR populations have significantly higher polygenic risk scores than IBS and TSI, p=1.5e-75, **Figure S9**A), polygenic scores for height across super populations show biased predictions. For example, the African populations sampled are genetically predicted to be considerably shorter than all Europeans and minimally taller than East Asians (Figure 4A), which contradicts empirical observations (with the exception of some indigenous pygmy/pygmoid populations)^77,78^. Additionally, polygenic risk scores for schizophrenia, while at a similar prevalence across populations where it has been well-studied^79^ and sharing significant genetic risk across populations^80^, shows considerably decreased scores in Africans compared to all other populations (Figure 4B). Lastly, the relative order of polygenic risk scores computed for type II diabetes across populations differs depending on whether the summary statistics are derived from a European-specific (Figure 4C) or multi-ethnic (Figure 4D) cohort.

**Figure 4.**
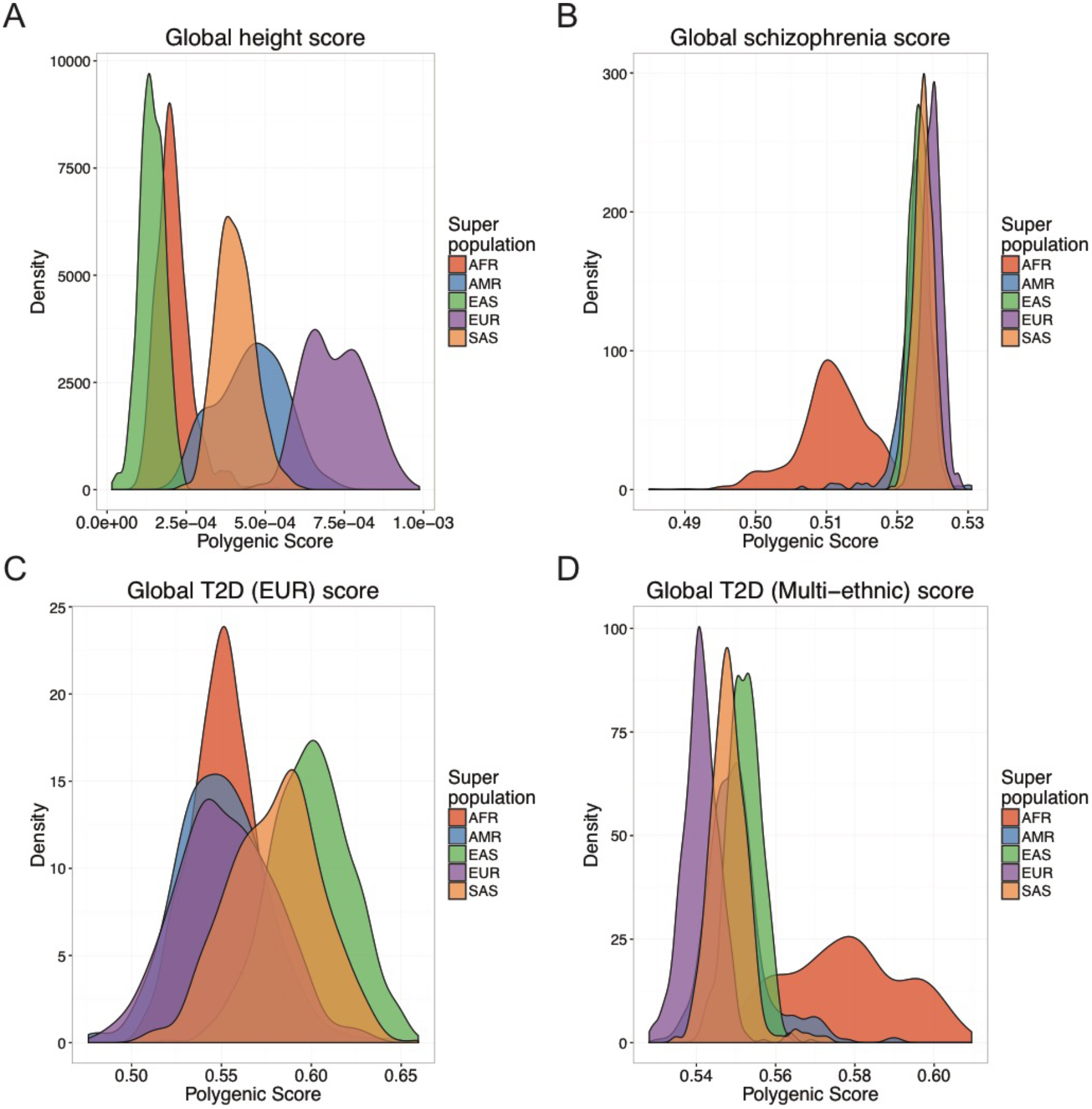
Biased genetic discoveries influence disease risk inferences. A-D) Inferred polygenic risk scores across individuals colored by population for: A) height based on summary statistics from ^9^. B) schizophrenia based on summary statistics from ^10^. C) type II diabetes summary statistics derived from a European cohort from ^74^. D) type II diabetes summary statistics derived from a multiethnic cohort from ^75^.

#### Ancestry-specific biases in polygenic risk score estimates

We performed coalescent simulations to determine how GWAS signals discovered in one ancestral case/control cohort (i.e. ‘single-ancestry’ GWAS) are expected to impact polygenic risk score estimates in other populations under neutrality using summary statistics (for details, see Methods). Briefly, we simulated variants according to a previously published demographic model inferred from Africans, East Asians, and Europeans^13^. We specified “causal” alleles and effect sizes randomly, such that each causal variant has evolved neutrally and has a mean effect of zero with the standard deviation equal to the global heritability divided by number of causal variants. We then computed the true polygenic risk for each individual as the product of the estimated effect sizes and genotypes, then standardized the scores across all individuals. We calculated the total liability as the sum of the genetic and random environmental contributions, then identified 10,000 European cases with the most extreme liabilities and 10,000 other European controls. We then computed Fisher’s exact tests with this European case-control cohort, then quantified inferred polygenic risk scores as the sum of the product of genotypes and log odds ratios for 10,000 samples per population not included in the GWAS.

In our simulations and consistent with realistic coalescent models, most variants are rare and population-specific; “causal” variants are sampled from the global site frequency spectrum, resulting in subtle differences in true polygenic risk across populations (**Figure S12**, Figure 5B-D). We mirrored standard practices for performing a GWAS and computing polygenic risk scores (see above and Methods). We find that the correlation between true and inferred polygenic risk is generally low (Figure 5A, **Figure S13**), consistent with limited variance explained by polygenic risk scores from GWAS of these cohort sizes for height (e.g. ~10% of variance explained for a cohort of size 183,727^81^) and schizophrenia (e.g. ~7% variance explained for a cohort of size 36,989 cases and 113,075 controls^10^). Low correlations in our simulations are most likely because common tag variants are a poor proxy for rare causal variants. As expected, correlations between true and inferred risk within populations are typically highest in the European population (i.e. the population in which variants were discovered, Figure 5A and **Figure S13**). Across all populations, the mean Spearman correlations between true and inferred polygenic risk increase with increasing heritability while the standard deviation of these correlations significantly decreases (p=0.05); however, there is considerable within-population heterogeneity resulting in high variation in scores across all populations. We find that in these neutral simulations, a polygenic risk score bias in essentially any direction is possible even when choosing the exact same causal variants, heritability, and varying only fixed effect size (i.e. inferred polygenic risk in Europeans can be higher, lower, or intermediate compared to true risk relative to East Asians or Africans, **Figure S12**, Figure 5B-D).

**Figure 5.**
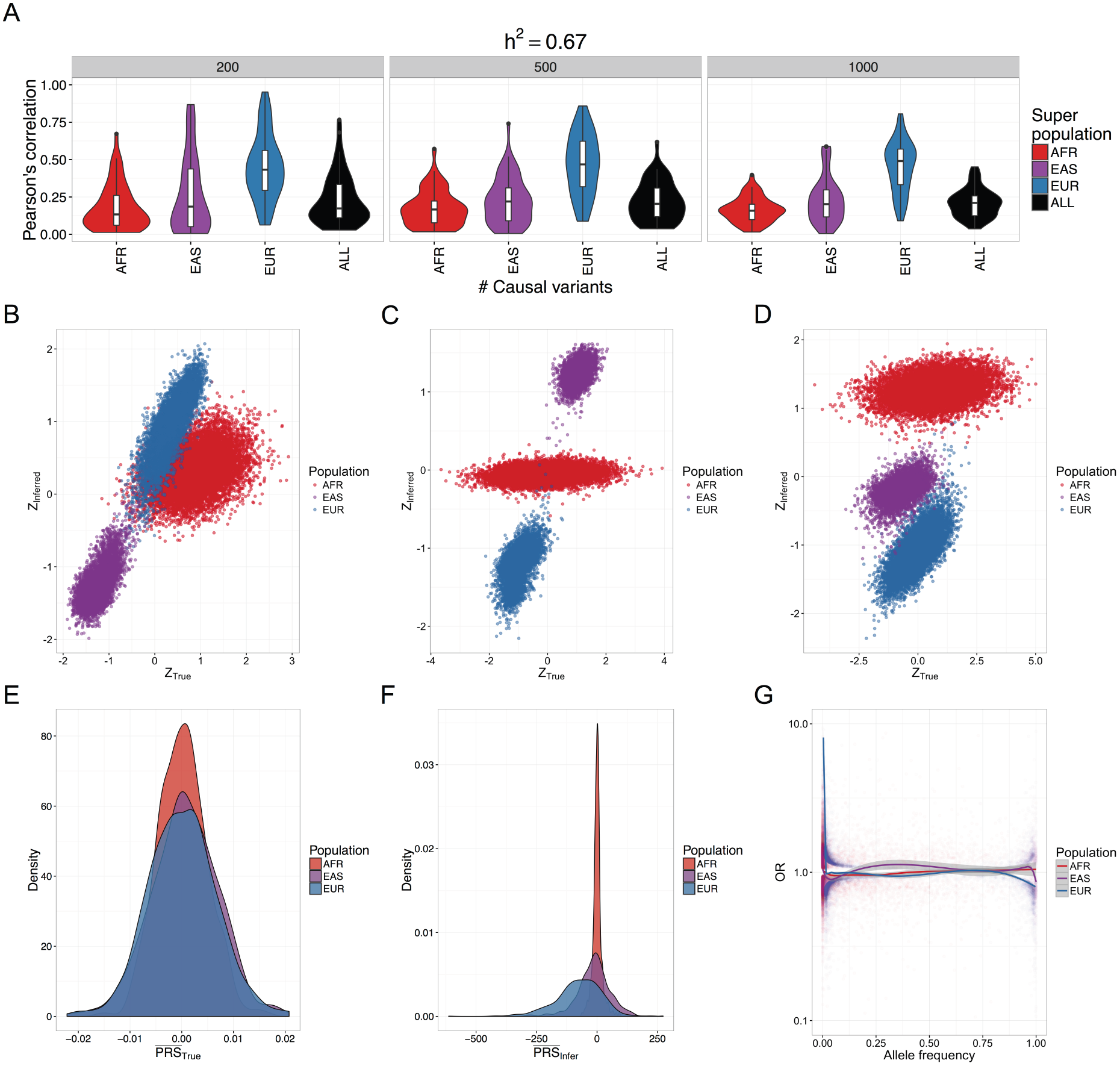
Coalescent simulation results for true vs inferred polygenic risk scores computed from GWAS summary statistics with 10,000 cases and 10,000 controls modeling European, East Asian, and African population history (demographic parameters from ^13^). A) Violin plots show Pearson’s correlation across 50 iterations per parameter set between true and inferred polygenic risk scores across differing genetic architectures, including *m*=200, 500, and 1,000 causal variants and h^2^=0.67. The “ALL” population correlations were performed on population mean-centered true and inferred polygenic risk scores. B-D) Standardized true versus inferred polygenic risk scores for 3 different coalescent simulations showing 10,000 randomly drawn samples from each population not included as cases or controls. E-F) The distribution for each population across 500 simulations with *m*=1000 causal variants and h^2^=0.67 of: E) unstandardized mean true polygenic risk and F) unstandardized mean inferred polygenic risk. G) Allele frequency versus inferred odds ratio for sites included in inferred polygenic risk scores for each population across 500 simulations, as in E-F).

While causal variants in our simulations are drawn from the global site frequency spectrum and are therefore mostly rare, inferred scores are derived specifically from common variants that are typically much more common in the study population than elsewhere (here Europeans with case/control MAF ≥ 0.01). Consequently, the distribution of mean true polygenic risk across simulation runs for each population are not significantly different (Figure 5E); however, inferred risk is considerably less than zero in Europeans (p=1.9e-54, 95% CI=[-84.3, -67.4]), slightly less than zero in East Asians (p=5.9e-5, 95% CI=[-19.1, -6.6]) and not significantly different from zero in Africans, with variance in risk scores decreasing with this trend (Figure 5F). The scale is orders of magnitude different between the true and inferred unstandardized scores, cautioning that while they are informative on a relative scale (Figure 5A and **Figure S11**), their absolute scale should not be overinterpreted. The inferred risk difference between populations is driven by the increased power to detect minor risk alleles rather than protective alleles in the study population^82^, given the differential selection of cases and controls in the liability threshold model. We demonstrate this empirically in these neutral simulations within the European population (Figure 5G), indicating that this phenomenon occurs even in the absence of population structure and when case and control cohort sizes are equal.

### Discussion

To date, GWAS have been performed opportunistically in primarily single-ancestry European cohorts, and an open question remains about their biomedical relevance for disease associations in other ancestries. As studies gain power by increasing sample sizes, effect size estimates become more precise and novel associations at lower frequencies are feasible. However, rare variants are largely population-private, and their effects are unlikely to transfer to new populations. Because linkage disequilibrium and allele frequencies vary across ancestries, effect size estimates from diverse cohorts are typically more precise than from single-ancestry cohorts (and often tempered)^5^, and the resolution of causal variant fine-mapping is considerably improved^75^. Across a range of genetic architectures, diverse cohorts provide the opportunity to reduce false positives. At the Mendelian end of the spectrum, for example, disentangling risk variants with incomplete penetrance from benign false positives and localizing functional effects in genes is much more feasible with large diverse population cohorts than possible with single-ancestry analyses^83,84^. Multiple false positive reports of pathogenic variants causing hypertrophic cardiomyopathy, a disease with relatively simple genomic architecture, have been returned to patients of African descent or unspecified ancestry that would have been prevented if even a small number of African American samples were included in control cohorts^85^. At the highly complex end of the polygenicity spectrum, we and others have shown that the utility of polygenic risk inferences and the heritable phenotypic variance explained in diverse populations is improved with more diverse cohorts^80,86^.

Standard single-ancestry GWAS typically apply linear mixed model approaches and/or incorporate principal components as covariates to control for confounding from population structure with primarily European-descent cohorts^1–3^. A key concern when including multiple diverse populations in a GWAS is that there is increasing likelihood of identifying false positive variants associated with disease that are driven by allele frequency differences across ancestries. However, previous studies have analyzed association data for diverse ancestries and replicated findings across ethnicities, assuaging these concerns^7,75,87^. In this study, we show that this ancestry stratification is not continuous along the genome: long tracts of ancestrally diverse populations present in admixed samples from the Americas are easily and accurately detected. Querying population substructure within these tracts recapitulates expected trends, e.g. European ancestry in African Americans primarily descends from northern Europeans in contrast to European ancestry from Hispanic/Latinos, which primarily descends from southern Europeans, as seen previously^49^. Additionally, population substructure follows a north-south cline in the Native component of Hispanic/Latinos, and the African component of admixed African descent populations in the Americas most closely resembles reference populations from Nigeria (albeit given the limited set of African populations from The 1000 Genomes Project). Admixture mapping has been successful at large sample sizes for identifying ancestry-specific genetic risk factors for disease^88^. Given the level of accuracy and subcontinental-resolution attained with local ancestry tracts in admixed populations, we emphasize the utility of a unified framework to jointly analyze genetic associations with local ancestry simultaneously^45^.

The transferability of GWAS is aided by the inclusion of diverse populations^89^. We have shown that European discovery biases in GWAS are recapitulated in local ancestry tracts in admixed samples. We have quantified GWAS study biases in ancestral populations and shown that GWAS variants are at lower frequency specifically within African and Native tracts and higher frequency in European tracts in admixed American populations. Imputation accuracy is also stratified across diverged ancestries, including across local ancestries in admixed populations. With decreased imputation accuracy especially on Native American tracts, there is decreased power for potential ancestry-specific associations. This differentially limits conclusions for GWAS in an admixed population in a two-pronged manner: the ability to capture variation and the power to estimate associations.

As GWAS scale to sample sizes on the order of hundreds of thousands to millions, genetic risk prediction accuracy at the individual level improves^90^. However, we show that the utility of polygenic risk scores computed using GWAS summary statistics are dependent on genetic similarity to the discovery cohort. BLUP risk prediction methods have been proposed to improve risk scores, but they require access to raw genetic data typically from very large datasets, are also dependent on LD structure in the study population, and only offer modest improvements in prediction accuracy^91^. Furthermore, polygenic risk scores contain a mix of true positives (which have the bias described above) and false positives in the training GWAS. False positives, being chance statistical fluctuations, do not have the same allele frequency bias and therefore unfortunately play an outsized role in applying a PRS in a new population.

We have demonstrated that polygenic risk computed from summary statistics in a single-ancestry cohort can be biased in essentially any direction across diverse populations simply as a result of genetic drift, limiting their interpretability; directional selection is expected to bias polygenic risk inferences even more. Because biases arise from genetic drift alone, we recommend: 1) avoiding interpretations from polygenic risk score differences extrapolated across populations, as these are likely confounded by latent population structure that is not properly corrected for with current methods, 2) mean-centering polygenic risk scores for each population, and 3) computing polygenic risk scores in populations with similar demographic histories as the study sample to ensure maximal predictive power. Further, additional methods that account for local ancestry in genetic risk prediction to incorporate different ancestral linkage disequilibrium and allele frequencies are needed. This study demonstrates the utility of disentangling ancestry tracts in recently admixed populations for inferring recent demographic history and identifying ancestry-stratified analytical biases; we also motivate the need to include more ancestrally diverse cohorts in GWAS to ensure that health disparities arising from genetic risk prediction do not become pervasive in individuals of admixed and non-European descent.

## Author contributions

ARM, CRG, RKW, CDB, MJD, EEK conceived of and designed the experiments. ARM and GLW performed the data analysis. SG, CDB, and MJD contributed analysis tools/materials. ARM wrote the manuscript with comments from CRG, RKW, GLW, MJD, SG, and EEK. All authors read and approved the final manuscript.

## Acknowledgments

We thank Suyash Shringarpure, Brian Maples, Andres Moreno-Estrada, Danny Park, Noah Zaitlen, Alexander Gusev, and Alkes Price for helpful discussions/feedback. We thank Verneri Antilla for providing GWAS summary statistics. We thank Jerome Kelleher for several conversations about msprime, providing example scripts, and implementing new simulation capabilities. This work was supported by funds from several grants; the National Human Genome Research Institute under award numbers U01HG009080 (EEK, CDB, CRG), U01HG007419 (CDB, CRG, GLW), U01HG007417 (EEK), U01HG005208 (MJD), T32HG000044 (CRG) and R01GM083606 (CDB), and the National Institute of General Medical Sciences under award number T32GM007790 (ARM) at the National Institute of Health; the Directorate of Mathematical and Physical Sciences award 1201234 (SG, CDB) at the National Science Foundation; the Canadian Institutes of Health Research through the Canada Research Chair program and operating grant MOP-136855 (SG), and a Sloan Research Fellowship (SG).

## Web Resources

Phased 1000 Genomes haplotypes: ftp://ftp-trace.ncbi.nih.gov/1000genomes/ftp/release/20130502/supporting/shapeit2_scaffolds/w_gs_gt_scaffolds/

Local ancestry calls: https://personal.broadinstitute.org/armartin/tgp_admixture/

Scripts for processing data and running simulations: https://github.com/armartin/ancestry_pipeline/

PCAmask software: https://sites.google.com/site/pcamask/dowload

Tracts software: https://github.com/sgravel/tracts

Msprime software: https://github.com/jeromekelleher/msprime

